# Decoding Arc Transcription: A Live-Cell Study of Stimulation Patterns and Transcriptional Output

**DOI:** 10.1101/2024.03.28.587245

**Authors:** Dong Wook Kim, Hyungseok C. Moon, Byung Hun Lee, Hye Yoon Park

## Abstract

Activity-regulated cytoskeleton-associated protein (Arc) plays a crucial role in synaptic plasticity, a process integral to learning and memory. Arc transcription is induced within a few minutes of stimulation, making it a useful marker for neuronal activity. However, the specifics of the neuronal activity that triggers Arc transcription remain unknown because it has not been possible to observe mRNA transcription in live cells in real time. Using a genetically encoded RNA indicator (GERI) mouse model that expresses endogenous Arc mRNA tagged with multiple GFPs, we investigated Arc transcriptional activity in response to various electrical stimulation patterns. In dissociated hippocampal neurons, we found that the pattern of stimulation significantly affects Arc transcription. Specifically, a 10 Hz burst stimulation induced the highest rate of Arc transcription. Concurrently, the amplitudes of nuclear calcium transients also reached their peak with 10 Hz stimulation, indicating a correlation between calcium concentration and transcription. However, our dual-color single-cell imaging revealed that there were no significant differences in calcium amplitudes between Arc-positive and Arc-negative neurons upon 10 Hz burst stimulation, suggesting the involvement of other factors in the induction of Arc transcription. Our live-cell RNA imaging provides a deeper insight into the complex regulation of transcription by activity patterns and calcium signaling pathways.

## Introduction

Learning and memory are complex cognitive functions that are critical for the survival and adaptation of an organism. A central biological mechanism underlying these functions is synaptic plasticity, the ability of synapses to change their strength and efficiency in response to varying levels of neuronal activity (Ho et al., 2011; Magee and Grienberger, 2020). This dynamic process is modulated by a complex set of molecular mechanisms. Among these, activity-dependent gene expression plays an indispensable role in the regulation of synaptic plasticity (Kandel, 2001; Flavell and Greenberg, 2008). The induction of specific genes following neuronal stimulation leads to the production of proteins that can restructure synaptic connections. As neuronal activity levels fluctuate, this gene expression adapts accordingly, fostering the creation of dynamic neural networks that shape in accordance with an organism’s experiences. Proteins produced as a result of this activity-dependent gene expression can augment synaptic strength, form new synaptic connections, or prune unnecessary ones, thereby sculpting the neural architecture that forms the basis of learning and memory.

The onset of activity-dependent gene expression is primarily characterized by the activation of immediate early genes (IEGs) (Sheng and Greenberg, 1990; Minatohara et al., 2016). These genes exhibit a rapid and transient response to a wide range of cellular stimuli, including neuronal activity. Most IEGs encode proteins that act as transcription factors, influencing the expression of other genes and modulating neuronal function and synaptic plasticity. A notable exception is the activity-regulated cytoskeleton-associated protein (Arc/Arg3.1), a well-characterized IEG. Unlike most IEGs, Arc is an effector protein that acts directly at synapses (Zhang and Bramham, 2020). Arc protein is involved in AMPA receptor endocytosis, changes in synaptic morphology, and regulation of actin dynamics, which collectively contribute to synaptic plasticity (Chowdhury et al., 2006; Nair et al., 2017; Zhang and Bramham, 2020). However, despite its well-established role, the effect of neuronal activity patterns on Arc transcriptional dynamics remains poorly understood, largely due to the challenge of observing Arc mRNA transcription in real time in living cells.

Previously, transcription has been studied either by isolating RNA after breaking up the cells, or by *in situ* hybridization, which can localize specific gene transcripts to individual cells (Vera et al., 2016). However, the former does not recapitulate the complex interactions and the spatial relationships between the cells, and the latter requires fixation of the sample, which loses the dynamic information of the process. Most live-cell imaging methods rely on the use of exogenous reporter genes, which typically do not contain all cis-regulatory elements, raising doubts about the physiological relevance of the results.

To decipher the neuronal activity pattern that triggers Arc transcription, we employed a genetically encoded RNA indicator (GERI) mouse model (Lee et al., 2022). This mouse model expresses endogenous Arc mRNA tagged with multiple GFPs, enabling real-time monitoring of Arc transcriptional dynamics in live neurons. In this study, we specifically focused on the effects of different electrical stimulation patterns on Arc transcription in dissociated hippocampal neurons. We also examined the link between Arc transcription and calcium transient amplitude, a factor implicated in numerous signaling pathways in synaptic plasticity. These results shed new light on the relationship between neuronal activity and transcriptional output in live neurons.

## Results

### Imaging Arc mRNA transcription in live neurons

To visualize transcription of Arc mRNA in live neurons, we utilized the specific binding of PP7 capsid protein (PCP) to the stem loop structure of PP7 binding site (PBS). We used the PCP×PBS GERI mouse model, which was generated by crossing the Arc-PBS knock-in (KI) mouse (Das et al., 2018) with the PCP-GFP transgenic mouse (Lee et al., 2022). In the Arc-PBS KI mouse, 24 repeats of the PBS are knocked in the 3’untranslated region (3’UTR) of Arc mRNA (Das et al., 2018) (Figure 1A, left). In the PCP-GFP transgenic mouse, tandem PCP fused with tandem GFP is expressed under the human synapsin promoter (Lee et al., 2022) (Figure 1A, right). We cultured hippocampal neurons from PCP×PBS mice carrying homozygous Arc-PBS and heterozygous PCP-GFP transgenes (Figure 1A). In these neurons, all endogenous Arc mRNAs are labeled with up to 48 GFPs with a relatively low PCP-GFP background (Lee et al., 2022) (Figure 1B).

**Figure 1.**
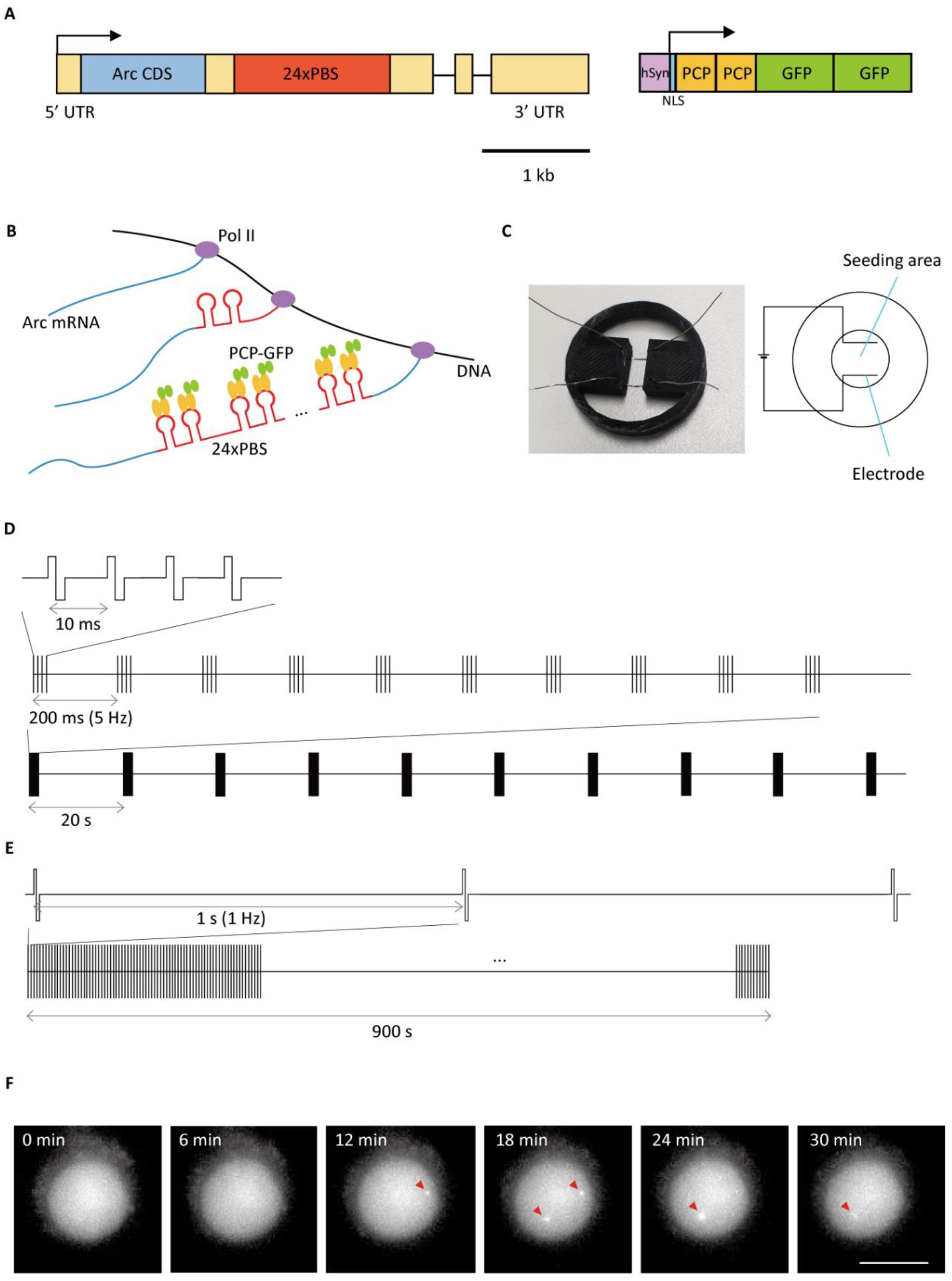
Arc-PBS tagging and electrical stimulation system for investigating Arc transcription in live neurons. **(A)** Schematic of the Arc-PBS gene with 24×PBS cassette inserted in the 3′ UTR region. Yellow boxes, UTR; black lines, introns; NLS, nuclear localization sequence. **(B)** Arc mRNAs labeled with 24×PBS repeats are transcribed from the template DNA, and PCP-GFP fusion proteins bind to the 24×PBS repeats. Pol II, RNA polymerase II. **(C)** Schematic of electrical stimulation system. **(D)** Schematic of theta burst stimulation (TBS). Each burst consists of 4 biphasic pulses with 2 ms duration and 10 ms intervals (top). Each train consists of 10 bursts with 200 ms intervals (middle). TBS consists of 10 trains with 20 s intervals (bottom). **(E)** Schematic of low frequency stimulation (LFS). LFS consists of 900 pulses with 1 s interval. **(F)** Time-lapse images of Arc transcription induced by TBS. Red arrowheads indicate transcription sites of Arc mRNA. Scale bar, 10 μm.

To stimulate neurons, we constructed a three-dimensional (3D) printed frame that holds two parallel platinum wires in close proximity to the neurons on a glass-bottom dish and applied electric fields (Figure 1C). First, we applied stimulation patterns known to induce long-term potentiation (LTP) and long-term depression (LTD). For LTP induction, we used theta burst stimulation (TBS) pattern. This pattern comprises of 4 pulses at 100 Hz in each burst, with a total of 10 bursts per train separated by intervals of 200 ms (Figure 1D). On the other hand, to induce LTD, we employed low frequency stimulation (LFS). This pattern consists of 900 pulses at 1 Hz (Balkowiec and Katz, 2002; Grover et al., 2009) (Figure 1E). Transcription of Arc mRNA was observed in live neurons after both TBS (Figure 1F and Movie 1) and LFS.

### Calcium activity and Arc transcription induced by TBS and LFS

Since electrical stimulation induces calcium activity and triggers transcription of immediate-early genes, we monitored calcium activity using a ratiometric calcium indicator, Fura Red. In this study, we focused on the effects of nuclear calcium transients on the induction of Arc transcription. Fluorescence signals representing nuclear calcium transients were recorded during stimulation. The region of interest (ROI) in each cell was selected based on the nuclear GFP signal due to the nuclear localization sequence (NLS) in the tandem PCP fused with tandem GFP construct. We obtained the fluorescence intensity ratio, *R* = *F*_405nm_/*F*_488nm_, by dividing the emission at 405 nm excitation by the emission at 488 nm excitation (Figure 2A and 2B). Figure 2C shows the average Δ*R*/*R* profile of the neurons stimulated by TBS (n = 19 cells from 3 independent neuron cultures). Each train of TBS elicited abrupt surges in calcium concentration. On the other hand, LFS resulted in an initial calcium spike followed by a steady calcium level throughout the stimulation period (Figure 2D; n = 13 cells from 3 independent neuron cultures).

**Figure 2.**
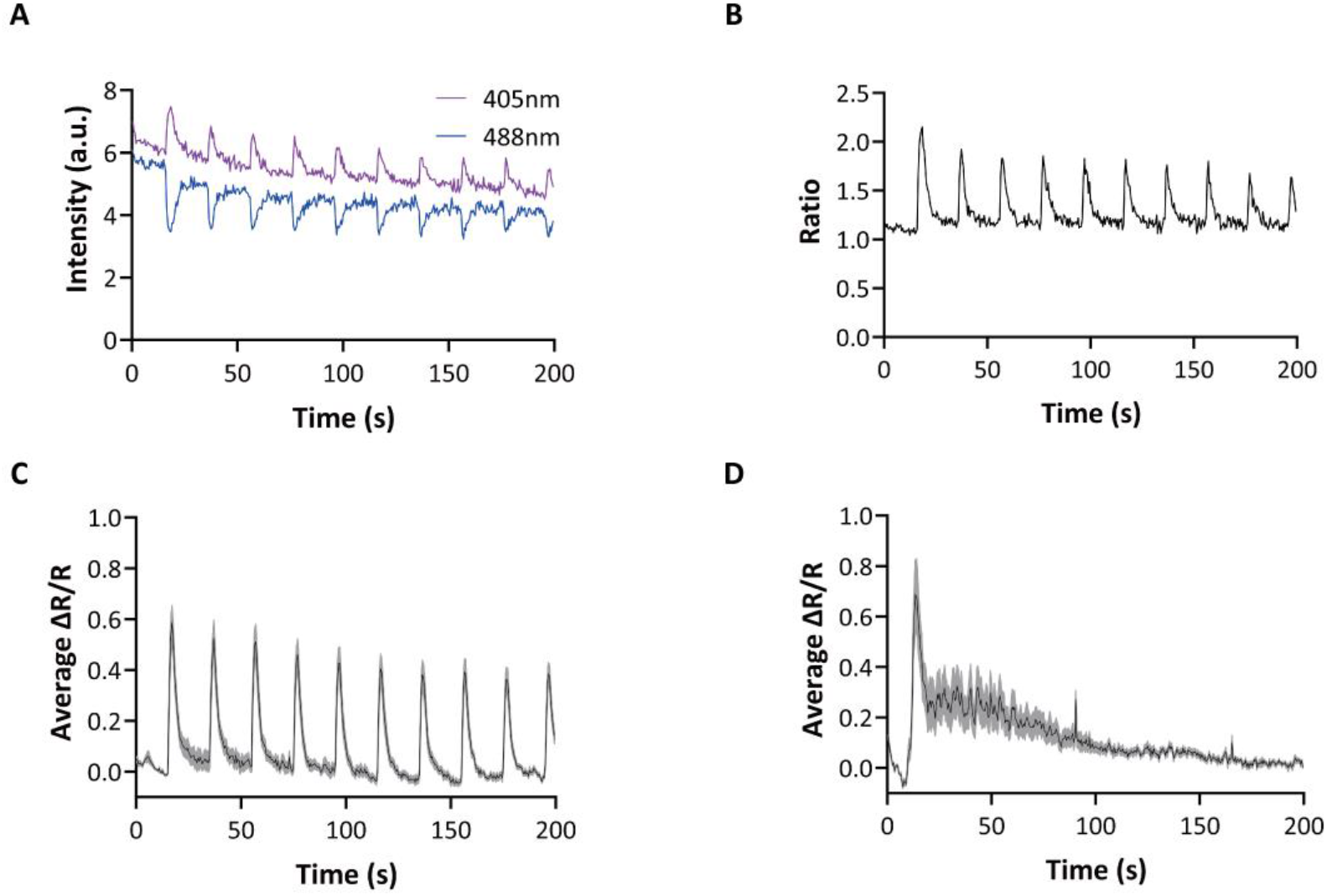
Calcium activity elicited by TBS and LFS. **(A)** Fluorescence intensities of Fura Red in a neuron during 10 Hz burst stimulation; *F*_405nm_ at 405 nm and *F*_488nm_ at 488 nm excitation. **(B)** Calcium intensity ratio calculated by *F*_405nm_/*F*_488nm_. **(C)** Average ΔR/R profile of the cells stimulated by TBS (n = 19 cells from 3 independent neuron cultures). **(D)** Average ΔR/R profile of the cells stimulated by LFS (n = 13 cells from 3 independent neuron cultures).

We then examined Arc mRNA transcription after TBS or LFS in live neurons. The fraction of neurons showing one or two Arc transcription sites was highest at 12-24 min after the onset of either stimulation (Figure 3A). While there exist transcribing cells in basal condition due to spontaneous activities (Das et al., 2018), we found that both TBS and LFS rapidly induced *de novo* Arc transcription within ∼ 20 min (Figure 3B). Because Arc transcription gradually decreased thereafter (Figure 3A), we focused on the transcriptional activity within 30 min of stimulation. The neurons that showed transcription sites during this period are hereafter referred to as Arc+ cells. The average fractions of Arc+ cells were 33 ± 3% after TBS and 27 ± 3% after LFS, which were significantly higher than 13 ± 2% without any stimulation (negative control, NC) (Figure 3C). The average onset time, the time to observe the first transcription site within 30 min, was ∼13 min for all three conditions (Figure 3D). The duration of Arc transcription on-time was also similar (16-18 min) for all three conditions (Figure 3E). Therefore, both TBS and LFS increased the probability of transcription activation, but not the onset time nor the duration of on-time.

**Figure 3.**
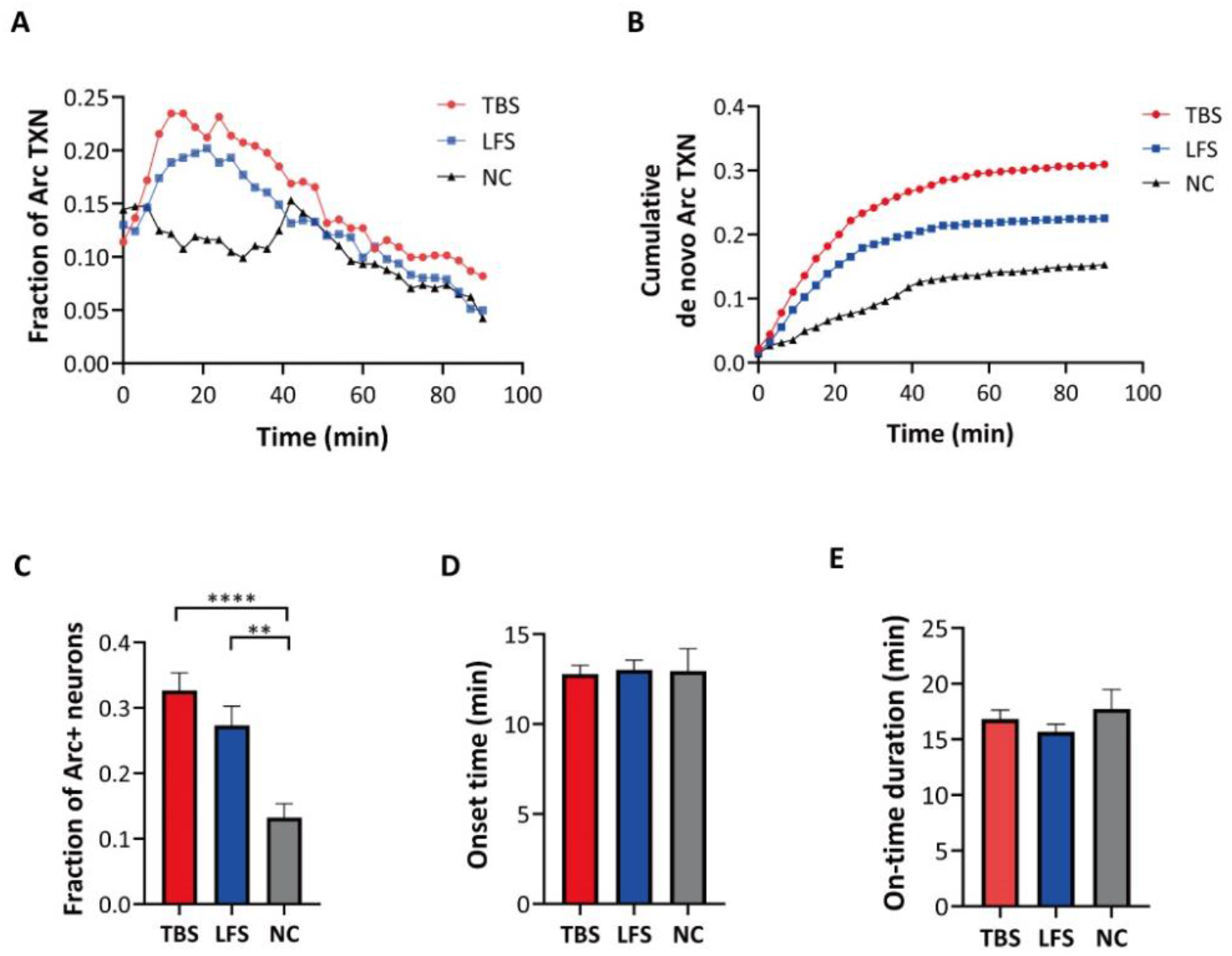
Transcription of Arc mRNA induced by TBS and LFS. **(A)** Time course of Arc transcription. Each point represents the fraction of cells that were transcribing Arc mRNA. **(B)** Time course of the cumulative fraction of *de novo* Arc transcription alleles. **(C)** Fraction of cells showing Arc transcription within 30 min of stimulation (n = 622 cells from 4 independent neuron cultures for TBS, n = 684 cells from 4 independent neuron cultures for LFS, n = 353 cells from 3 independent neuron cultures for NC, P < 0.0001 for one-way ANOVA test, **P < 0.01, ****P < 0.0001, *post hoc* Tukey’s t-tests). Imaging and stimulation started simultaneously at t = 0. TBS (10 stimulations of 5-Hz bursts) was given for 200 s and LFS (900 pulses at 1 Hz) was given for 900 s. **(D)** Mean onset time of Arc transcription (n = 622 cells for TBS, n = 684 cells for LFS, n = 353 cells for NC. P = 0.9456 for one-way ANOVA test). **(E)** Mean on-time duration of Arc transcription (n = 622 cells for TBS, n = 684 cells for LFS, n = 353 cells for NC. P = 0.3855 for one-way ANOVA test). Error bars indicate standard error of mean (SEM).

### Effect of electrical burst frequency on Arc transcription

Next, we sought to determine whether a specific frequency of the electrical bursts could significantly affect Arc transcription. We varied the interval between each burst (Figure 1D, middle) from 50 ms to 1 s, while keeping the other parameters of the burst stimulation constant. The average Δ*R*/*R* profiles at 1, 5, 10, 15, and 20 Hz burst stimulation (Figure 4A-E) indicate that 10 Hz burst induced significantly higher calcium amplitude than the other frequencies at each stimulation train (Figure 4F).

**Figure 4.**
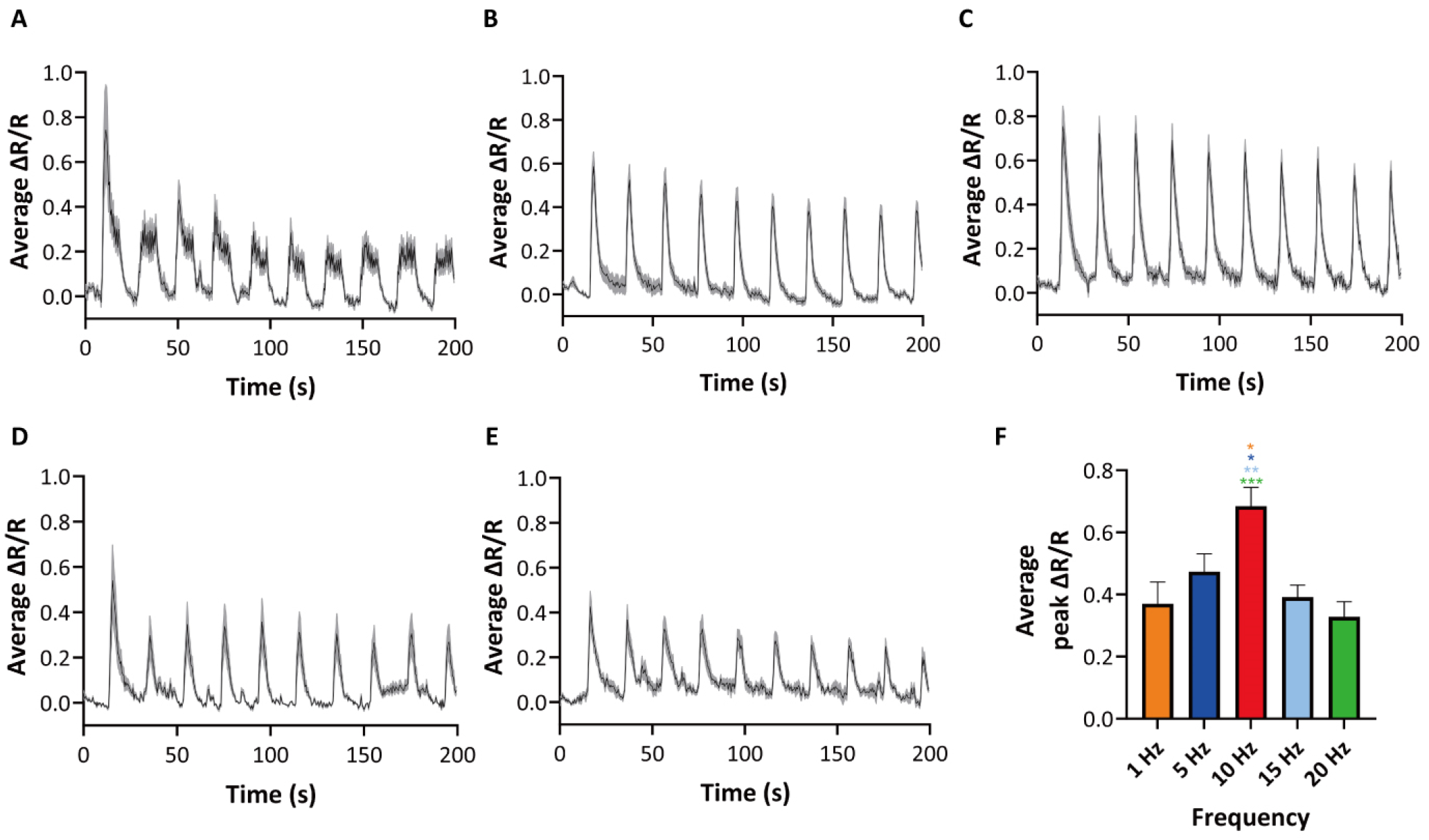
Calcium activity elicited by electrical burst stimulation with various burst frequencies. **(A-E)** Average ΔR/R profiles during burst stimulation at the frequency of 1 Hz (A), 5 Hz (B), 10 Hz (C), 15 Hz (D), and 20 Hz (E). **(F)** Average peak value of ΔR/R at each frequency (n = 7-19 cells, from 3 independent neuron cultures for each experiment, P = 0.0002 for one-way ANOVA test, *P < 0.05, **P < 0.01, ***P < 0.001, *post hoc* Tukey’s t-tests). Error bars indicate SEM.

We then examined the fractions of Arc transcribing cells (Figure 5A) and cumulative *de novo* Arc transcription alleles (Figure 5B) after the onset of each stimulation pattern. We found that 10 Hz burst stimulation also resulted in the highest Arc transcription rate compared to the other frequencies (Figure 5A and B). All tested burst frequencies (1-20 Hz) induced Arc transcription in a greater proportion of neurons than the negative control (no stimulation). Notably, we found a significantly higher proportion of Arc+ neurons after 10 Hz burst stimulation than any other stimulation frequency (Figure 5C). However, the average onset time of Arc transcription was 12-14 min for all 6 conditions (Figure 5D). These data suggest that there exists an optimal frequency for inducing Arc transcription. The results also highlight a correlated response in calcium amplitude and Arc transcription rate, both modulated by stimulation patterns.

**Figure 5.**
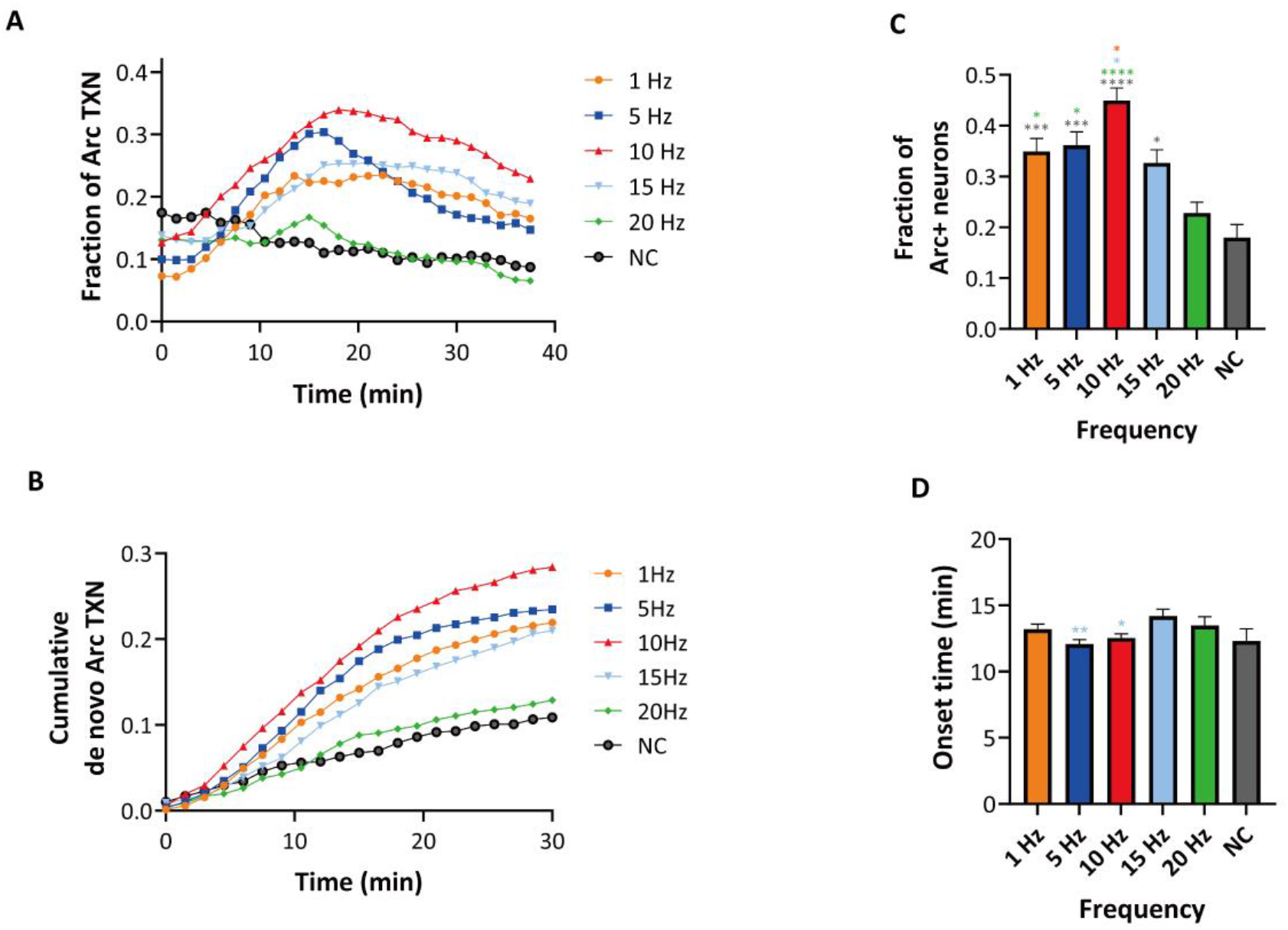
Transcription of Arc mRNA induced by the electrical burst stimulation with various burst frequencies. **(A)** Time course of Arc transcription. Each point represents the fraction of cells that were transcribing Arc mRNA at each time. **(B)** Cumulative fraction of *de novo* Arc transcription alleles. **(C)** Fraction of cells that transcribed Arc mRNA within 30 min after the onset of stimulation (n = 436-1096 cells, from 3-6 independent neural cultures, P < 0.0001 for one-way ANOVA test, *P < 0.05, ***P < 0.001, ****P < 0.0001, *post hoc* Tukey’s t-tests). **(D)** Mean onset time of Arc transcription (n = 436-1096 cells, P = 0.0086 for one-way ANOVA test, *P < 0.05, **P < 0.01, *post hoc* Tukey’s t-tests). Error bars indicate SEM.

### Effect of CREB phosphorylation on Arc transcription

Given the similar trends observed in calcium amplitude and Arc transcription at the population level, we sought to determine whether the amplitude of the calcium transient governs Arc transcription at the single cell level. To do this, we imaged both Arc transcription and calcium activity in the same cells (Figure 6A), whereas in the previous experiments we imaged calcium and transcriptional activity separately. We began with recording baseline images of Arc mRNA in a neuron. Next, we imaged calcium signals of the neuron using Fura Red during 10 Hz burst stimulation. Lastly, we proceeded to image Arc mRNA in the neuron every 1 min after stimulation. Figure 6B and 6C show time-lapse images of a representative Arc+ and Arc-cell, respectively. The average Δ*R*/*R* profiles of the Arc+ and Arc-neurons are shown in Figure 6D and 6E, respectively. The average peak Δ*R*/*R* values were similar for Arc+ and Arc-neurons (Figure 6F). Therefore, we did not find a significant relationship between the calcium amplitude and Arc transcription at the single cell level.

**Figure 6.**
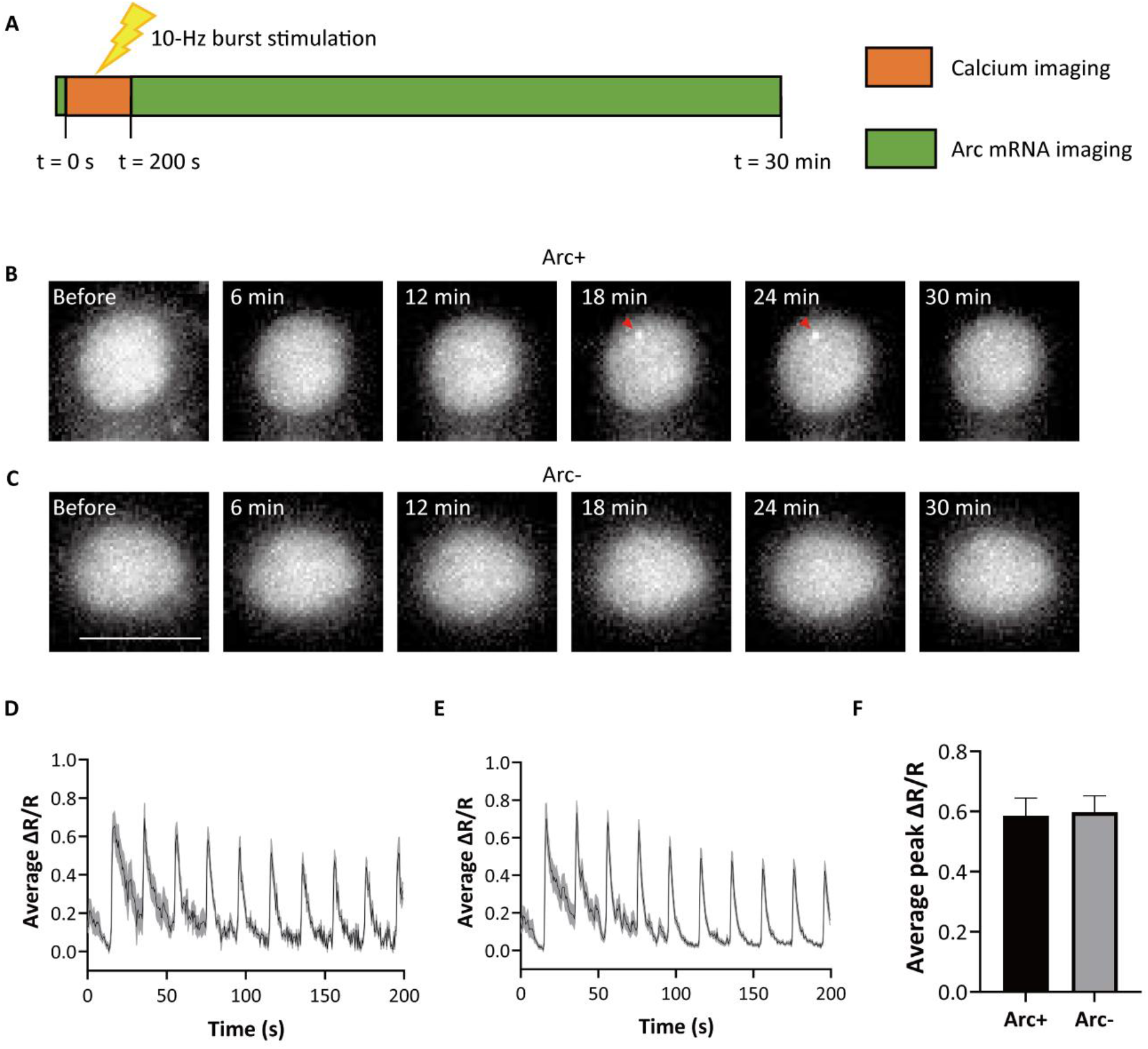
Calcium activity in Arc+ and Arc-cells. **(A)** Experimental scheme to record calcium activity and Arc transcription. **(B)** Time-lapse images of an Arc+ cell. Red arrowheads indicate transcription sites of Arc mRNA. **(C)** Time-lapse images of an Arc-cell. **(D)** Average ΔR/R profile of Arc+ cells (n = 10 cells from 3 independent neuron cultures). **(E)** Average ΔR/R profile of Arc-cells (n = 17 cells from 3 independent neuron cultures). **(F)** Average peak ΔR/R values of the Arc+ and Arc-cells. Scale bars, 10 μm. Error bars indicate SEM.

Next, we examined another key component of the signaling pathway – the cAMP response element-binding protein (CREB). CREB is a well-characterized transcription factor that is regulated by calcium activities. Calcium activity triggers a signaling cascade that leads to rapid phosphorylation of CREB and its binding to the cAMP response element (CRE) present in the promoter of many activity-regulated genes. Once bound to CRE, phosphorylated CREB (pCREB) recruits other proteins to form a transcription complex, facilitating initiation of target gene transcription (Bito et al., 1996; Hardingham et al., 2001; West et al., 2001). To detect any differences in CREB expression levels between Arc+ and Arc-neurons, we performed immunofluorescence (IF) and RNA fluorescence *in situ* hybridization (FISH) on neurons that were fixed after 10 Hz burst stimulation. Figure 7A shows a representative image of Arc-(left) and Arc+ (right) neurons expressing similar levels of CREB (magenta). Neurons were sorted into Arc+ and Arc-cells by the presence of one or two bright spots labeled with FISH probes (green) targeting the Arc coding sequence (Table 1) in the nucleus (blue). There was no significant difference in the mean intensity of CREB signal between Arc+ cells with one or two Arc transcription sites and Arc-cells (Figure 7B and C). There was no correlation between the CREB signal and the intensity of Arc transcription sites (Figure 7D). However, as shown in the representative image in Figure 7E, Arc+ cells exhibited higher levels of pCREB than Arc-cells (Figure 7F). Also, cells with two Arc transcription sites exhibited higher levels of pCREB than the cells with one Arc transcription site or Arc-cells (Figure 7G). However, there was no correlation between pCREB intensity and Arc transcription site intensity (Figure 7H). These results suggest that the on-rate of Arc transcription is correlated with the phosphorylation rate of CREB rather than the total CREB expression level.

**Figure 7.**
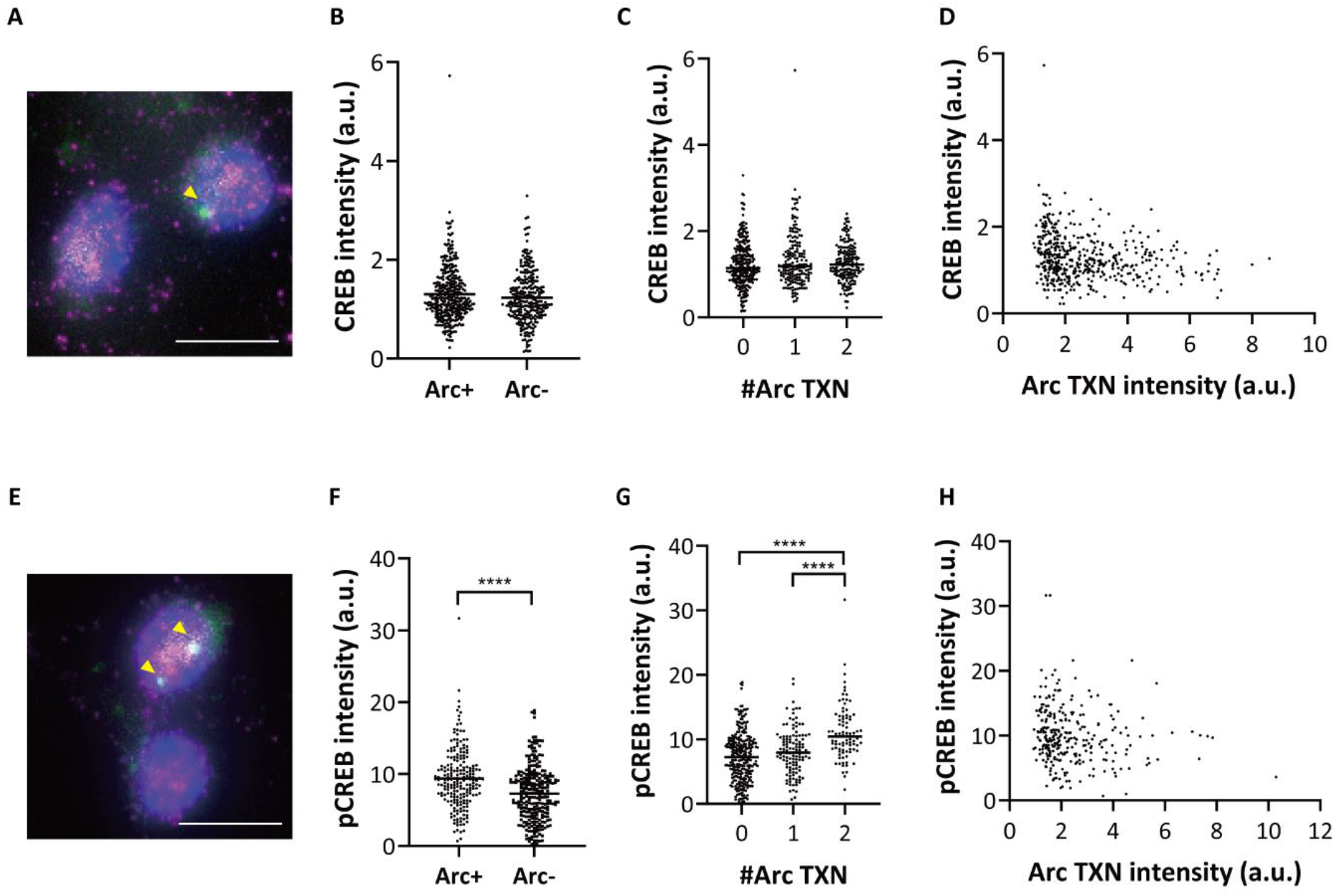
CREB and pCREB levels in Arc+ and Arc-cells. **(A, E)** Representative multichannel IF-smFISH image of CREB (A) or pCREB (E) protein (magenta), Arc mRNA (green), and DAPI (blue). Yellow arrowheads indicate transcription sites of Arc mRNA. **(B)** CREB intensity of Arc+ and Arc-cells (n = 346 for Arc+ cells, n = 279 for Arc-cells). **(C)** CREB intensity in neurons with 0, 1, and 2 Arc transcription sites (P = 0.1264 for one-way ANOVA test). **(D)** Scatter plot of CREB intensity and Arc transcription site intensity (n = 346 cells, r^2^ = 0.04469). **(F)** pCREB intensity of Arc+ and Arc-cells (n = 210 for Arc+ cells, n = 269 for Arc-cells, ****P < 0.0001, two-tailed t-test). **(G)** pCREB intensity and the number of Arc transcription allele (P < 0.0001 for one-way ANOVA test, ****P < 0.0001, *post hoc* Tukey’s t-tests). **(H)** Scatter plot of pCREB intensity and Arc transcription site intensity (n = 269 cells, r^2^ = 0.01987). Scale bars, 10 μm.

**Table 1.**
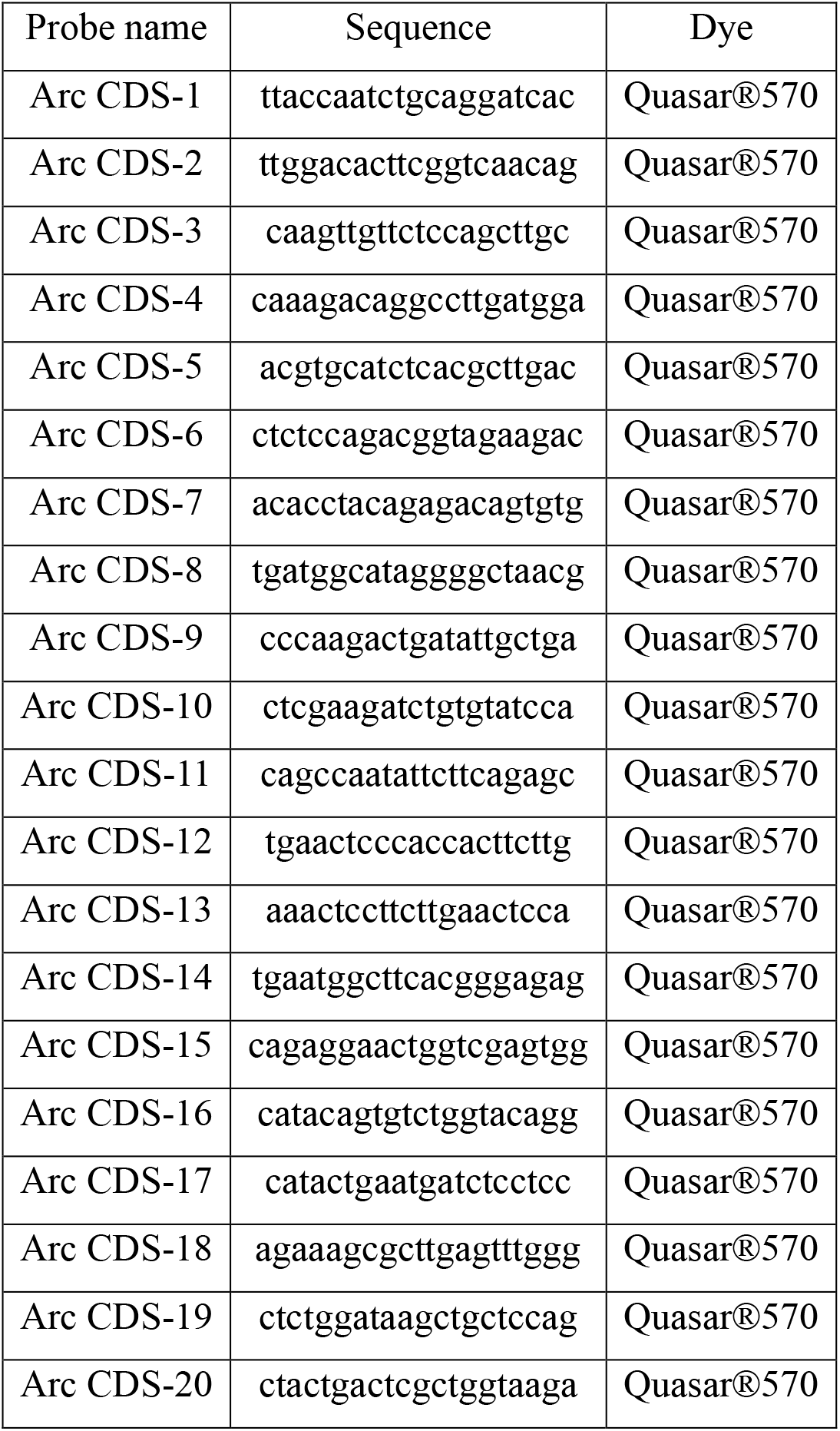
Sequence of the probes used for FISH against Arc coding sequence. Probe sets were modified at both 5′ and 3′ ends for conjugation to Quasar® 570 dyes.

## Discussion

In this study, we investigated the transcriptional response of neurons following diverse patterns of electrical stimulation. Using dissociated hippocampal neurons cultured from the GERI mouse model (Lee et al., 2022), we monitored transcriptional activity of endogenous Arc gene in live neurons. This approach allowed us to analyze hundreds of individual cells, yielding highly quantitative and statistical information about Arc transcription.

The role of Arc in synaptic plasticity and long-term memory formation has been the subject of extensive research. Several studies have demonstrated the role of Arc in the LTP induction and maintenance, as evidenced by impaired LTP in an Arc knock-out (KO) animal (Plath et al., 2006) or after infusion of Arc antisense oligodeoxynucleotides (Guzowski et al., 2000; Messaoudi et al., 2007). Interestingly, recent research has presented conflicting results, suggesting that LTP is unaffected in Arc KO animals (Kyrke-Smith et al., 2021). While uncertainty remains regarding the precise role of Arc in LTP, it has been established that Arc is crucial for LTD through the endocytosis of AMPAR receptors (Chowdhury et al., 2006; Park et al., 2008; Waung et al., 2008), implicating Arc in the regulation of Hebbian synaptic plasticity. Furthermore, evidence supports the idea that Arc plays a role in stabilizing synaptic responses, thereby influencing homeostatic plasticity (Shepherd et al., 2006; Gao et al., 2010; Peebles et al., 2010).

Since Arc is involved in multiple forms of synaptic plasticity, we first investigated whether the stimulation patterns known to induce either LTP or LTD could trigger Arc transcription in dissociated neurons. High frequency stimulation (HFS) and theta burst stimulation (TBS) are known to induce LTP (Bliss and Lomo, 1973; Larson et al., 1986; Capocchi et al., 1992), whereas low frequency stimulation (LFS) typically results in LTD (Dudek and Bear, 1992). Past studies indicated that while HFS or TBS could trigger Arc transcription, LFS fails to do so in the dentate gyrus *in vivo* (Link et al., 1995; Lyford et al., 1995; Steward et al., 1998; Waltereit et al., 2001). In the CA1 region, it was shown that LTP- and LTD-inducing stimulations lead to opposite changes in Arc mRNA levels *in vivo* (Yilmaz-Rastoder et al., 2011). The authors applied paired-pulse stimulation at 0.5 Hz to dorsal CA3 commissural fibers and observed LTD induction in CA1 neurons. However, Arc mRNA levels initially decreased and then displayed a transient increase after 70 min relative to control levels.

Our study, focusing on dissociated hippocampal neurons, found an increase in Arc transcription rates in response to both TBS and LFS. There was a significant surge in the number of neurons showing Arc transcription sites within 30 min of either TBS or LFS stimulation, compared to the negative control (Figure 3). We speculate that *de novo* Arc transcription might be necessary for LTD in cultured neurons due to low Arc levels in the culture preparation, whereas neurons *in vivo* might utilize pre-existing Arc mRNAs for rapid Arc translation (Wilkerson et al., 2018). Another possibility is that LFS triggers rapid degradation of Arc mRNA through nonsense-mediated mRNA decay (NMD) pathway, leading to a decrease or maintenance of total Arc mRNA levels despite *de novo* transcription (Yilmaz-Rastoder et al., 2011).

Next, we investigated how Arc transcription and calcium amplitude are modulated by different stimulation patterns. Interestingly, we found that the proportion of Arc+ cells varied with different stimulation patterns, with the highest Arc transcription rate observed at 10 Hz burst stimulation (Figure 5). This frequency falls within the theta oscillation range in rodents (6-10 Hz) (Buzsaki et al., 2013). This result is also in accordance with a previous study that found higher theta-burst activity in Arc+ cells *in vivo* (Lee et al., 2022). The highest Arc transcription and calcium amplitude at this specific burst frequency could be attributed to the cumulative effect of both excitatory and inhibitory influences from repeated burst stimulation (Larson and Munkacsy, 2015).

Our observations suggesting a correlation between Arc transcription and nuclear calcium concentration led us to hypothesize a potential difference in calcium amplitude between Arc+ and Arc-cells. However, when stimulated at the optimal burst frequency of 10 Hz, our results showed no distinguishable difference in the amplitude of calcium transients between Arc+ and Arc-cells (Figure 6F). This outcome suggests that factors downstream of calcium transients could also play a substantial role in inducing Arc transcription. One notable candidate is the CREB protein, which is known for its role in activity-dependent gene expression (Bito et al., 1996; Mayr and Montminy, 2001; Kornhauser et al., 2002; Lonze and Ginty, 2002). Upon calcium influx, CREB gets phosphorylated, binds to the synaptic activity-responsive element (SARE) in the enhancer region of the Arc gene, and triggers Arc transcription (Kawashima et al., 2009). We found significantly higher levels of pCREB but not total CREB expression levels in Arc+ neurons compared to Arc-neurons (Figure 7). These results suggest that the activation of CREB phosphorylation at optimal calcium levels further modulates Arc transcription. While some studies have shown that nuclear calcium transients regulate CREB phosphorylation and IEG transcription (Hardingham et al., 2001; Yu et al., 2017), other studies have suggested the importance of submembrane and cytosolic calcium transients to activate other intracellular pathways to initiate IEG transcription (Deisseroth et al., 1996; Nonaka et al., 2014; Colgan et al., 2023). In this study, we focused on the nuclear calcium signaling, but the effects of the calcium transients in other parts of the neurons could also be investigated in the future.

While we used hippocampal neural culture to observe the transcriptional response of individual live neurons, analysis on expression levels and spatial dynamics of Arc mRNA could be extended to various scales and regions using the Arc-PBS knock-in mouse model (Das et al., 2018). One of the recent studies have analyzed the transcriptional dynamics of single Arc mRNA in a time-resolved manner (Choi et al., 2022). By tracking Arc mRNA particles in the dendrites, other recent studies have revealed localization and transport dynamics of Arc mRNA (Das et al., 2018; Das et al., 2023). Compared to the field stimulation used in our study, synaptic stimulation in acute brain slices showed detailed aspects of Arc transcription *ex vivo* (Lituma et al., 2022). Since GERI imaging offers sensitive detection of pre-mRNA synthesis in individual neurons both in culture and *in vivo* (Lee et al., 2022), future studies using GERI in conjunction with electrophysiology or voltage imaging could provide important insights into the relationship between Arc transcription and electrical activity at the single-cell level.

In conclusion, our findings have provided insights into the dynamic nature of Arc transcription in response to different neuronal stimulation patterns. We have established that Arc transcription is sensitive to the type of stimulation and the amplitude of calcium transients, and is further regulated by downstream factors such as CREB phosphorylation. These results highlight the complexity of the mechanisms underlying activity-dependent gene expression and its potential implications for long-term synaptic plasticity. Although our study was carried out using dissociated neurons in a controlled environment, the principles revealed could provide a foundation for further studies investigating the relationship between neuronal activity and gene expression *in vivo*. Future research could benefit from exploring different mRNA species (Park et al., 2014; Lee et al., 2022) and brain regions in live mouse models, advancing our understanding of the intricate interplay between neuronal activity, gene expression, and synaptic plasticity.

## Materials and methods

### Hippocampal neuron preparation

All animal care and experiments were performed according to protocols approved by the Institutional Animal Care and Use Committee of Seoul National University. The day before neuronal culture, a glass-bottom dish (SPL, 100350) was coated with poly-D-lysine hydrobromide (Sigma-Aldrich P7886). The next day, hippocampi were dissected from the brains of 1-day-old Arc-PBS homozygous and PCP-GFP heterozygous mouse pups. The hippocampi were digested with 2.5% trypsin (Gibco, 15090046), plated onto the poly-D-lysine (PDL)-coated glass-bottom dishes at a concentration of 420,000 cells/mL, and incubated at 37 °C and 5% CO_2_ for ∼4 hours until they adhered to the bottom of the dishes. Then, B27 media consisting of 1× B-27 supplement (Gibco, 17504044), 1× GlutaMax supplement (Gibco, 35050079), and 0.1 mg/mL Primocin (InvivoGen, ant-pm-1) in Neurobasal-A media (Gibco, 10888022) was added to the dishes. Neurons were imaged between 16 and 18 days *in vitro* (DIV).

### Loading cells with calcium indicator dye

Fura Red ratiometric calcium indicator (Invitrogen #F3021) was used to monitor the intracellular calcium concentration of neurons. Prior to imaging, culture media were replaced with HEPES buffered saline (HBS) containing 119 mM NaCl, 5 mM KCl, 2 mM CaCl_2_, 2 mM MgCl_2_, 30 mM D-glucose, and 20 mM HEPES (Gibco #15630-080) at pH 7.4. Neurons were incubated in HBS with 4 μM Fura Red and 0.04% Pluronic® F-127 (Invitrogen #P3000MP) for 30 min at 37℃. Neurons were washed twice with HBS and incubated for another 30 min at 37℃ for de-esterification.

### Electrical stimulation

Electrical stimulation was delivered using a three-dimensional printed polylactic acid (PLA, Ultimaker) frame, which was designed to fit to the glass-bottom dish and hold two platinum wires close to the glass bottom. Each platinum wire was placed parallel to each other at a distance of 7.4 mm. Stimulation patterns were generated by an isolated pulse stimulator (A-M Systems, 2100) using trigger signals from the Arduino Uno. A single pulse was a biphasic pulse of 2 ms duration and 3.7 V amplitude. Low frequency stimulation (LFS) consisted of 900 pulses at a frequency of 1 Hz. Burst simulation consisted of 10 stimulations with an interval of 20 s, and each stimulation consisted of 10 bursts of various frequencies. Each burst consisted of 4 pulses at a frequency of 100 Hz.

### Live-cell imaging

To monitor Arc mRNA transcription, neurons were imaged by using an inverted microscope (Olympus IX83) equipped with an electron multiplying charge-coupled device (EMCCD) camera (Andor, iXon Ultra 888), 60 ×/1.3 NA silicon immersion objective (Olympus, UPLANSAPO60X), SOLA light-emitting diode (LED) Light Engine (Lumencor), motorized stage (Marzhauser), and Chamlide TC top stage incubator system (Live Cell Instruments). GFP signal was monitored with an ET-EGFP filter cube (Chroma, 49002). Z-stack images were acquired at 0.5 μm intervals. Neurons were imaged at 3-min intervals for TBS/LFS experiments and at 90-s intervals for frequency change experiments. For each experiment, neurons were imaged in the baseline condition before stimulation.

For real-time recording of intracellular calcium concentration, neurons were imaged using an inverted microscope (Olympus IX73) equipped with two EMCCD cameras (Andor, iXon Ultra 897), 60×/1.3 NA silicon immersion objective (Olympus, UPLANSAPO60X), MS-2000-500 XYZ automated stage (ASI), and Chamlide TC top stage incubator system (Live Cell Instruments). Calcium-bound Fura Red signals were obtained by excitation with a 405 nm diode laser (Cobolt), and calcium-free signals were obtained with a 488 nm diode laser (Cobolt). Each excitation laser was applied sequentially with an exposure time of 20 ms. Both emission signals were filtered with a 635/60 bandpass filter (Chroma). Calcium signals were recorded at 500 ms intervals. To compare the calcium signal between Arc+ and Arc-neurons, Arc mRNA signals were also observed immediately after calcium imaging. A 488 nm diode laser (Cobolt) was used to visualize Arc mRNA, and the fluorescence emission was filtered with a 525/50 bandpass filter (Chroma). Arc images were acquired at 60-s intervals.

### Arc transcription analysis

Imaging processing was performed using a similar approach described previously (Choi et al., 2022). Acquired z-stack images were projected to the maximum intensity profile at each time point. The mean intensity of a background region without cells was subtracted from the max-projected images. Transcription sites of Arc mRNA were tracked manually. When comparing Arc transcription rate between stimulation patterns, transcription sites activated before stimulation were excluded from the positive cell count. Cells showing transcription sites within 30 min of stimulation were considered positive cells. Time points at which transcription was initiated were also recorded, averaged across cells, and compared between stimulation patterns.

### Calcium signal analysis

We corrected shading and background variations in the time-lapse images by using the BaSiC plugin (Peng et al., 2017) and performed exponential bleach correction and background subtraction using Fiji (Schindelin et al., 2012). We selected cells expressing nuclear GFP signals and segmented the nuclei of the cells as ROIs. The calcium intensity ratio *R* was calculated by *F_405nm_*/*F_488nm_*. The initial baseline of calcium intensity ratio *R_0_* was calculated by averaging the bottom 50% of the first 50 frames, and the final baseline of the calcium intensity was calculated in the same way with the last 50 frames. For calcium intensity in Figure 6, the baseline was selected in the bottom 10% of the first or last 100 frames. *R* was adjusted by linearly fitting the final baseline to the initial baseline. Δ*R*/*R* was calculated as (*R*-*R*_0_)/*R*_0_. The calcium signal profile was obtained by averaging the Δ*R*/*R* of cells at each time point. The intensity of the peak signal elicited by each stimulation train was averaged per cell and compared between stimulation patterns.

### IF-FISH

Neurons were stimulated with 10-Hz burst stimulation and fixed with 4% PFA (Electron Microscopy Sciences #15714) on ice 15 min after the onset of stimulation. After fixation, neurons were washed in phosphate buffered saline with 1 mM MgCl_2_ and 1 mM CaCl_2_ (PBS-MC) and quenched in PBS-MC with 50 mM glycine. Then neurons were permeabilized with 0.1% Triton X-100 on ice and washed 3 times with PBS-MC. Neurons were incubated with 10% formamide, 2× SSC, and 0.5% BSA (Sigma, A7906) in RNase-free water for 30 min at room temperature for pre-hybridization and blocking. Neurons were hybridized with primary antibodies targeting CREB (Cell Signaling, 9197S) and pCREB (Sigma, 06-519), FISH probes targeting 20 sites in the open reading frame of the Arc gene (Stellaris; sequences in Table 1), 1 mg/ml tRNA (Sigma, R1753), 10% dextran sulfate (Sigma, D8906), 0.2 mg/ml BSA (Roche, 10711454001), 2× SSC, 2 mM vanadyl ribonucleoside complex (Sigma, R3380), and 10 U/ml SUPERase-In (Invitrogen, AM2694) in RNase-free water at 37℃ for 3 hr. After hybridization, neurons were washed twice with 10% Formamide and 0.5% BSA in 2× SSC and incubated twice with goat anti-rabbit secondary antibody labeled with Alexa 647 (Invitrogen, A21245), 10% Formamide, and 0.5% BSA in 2× SSC at 37℃ for 20 min each time. Neurons were washed 4 times with 2× SSC and stained with 0.5 µg/ml DAPI (Thermo Scientific, 62248) in 2× SSC for 1 min. Neurons were washed with PBS-MC for 10 min and mounted with Prolong Gold (Invitrogen, P10144).

### Arc-CREB analysis

After IF-FISH, neurons were imaged using an inverted microscope (Olympus IX83) equipped with the same devices described above. Arc-CDS probe signal was monitored with an ET-Cy3/TRITC filter cube (Chroma, 49004). The CREB/pCREB signal was monitored with an ET-Cy5 filter cube (Chroma, 49006). DAPI signal was monitored using an ET-DAPI filter cube (Chroma, 49000). The GFP signal was monitored using an ET-EGFP filter cube (Chroma, 49002). Z-stack images were acquired at 0.5 μm intervals. After multi-channel imaging, nuclear ROIs were selected based on DAPI and GFP signals. The number of Arc transcription sites in each cell was counted manually, and the mean CREB/pCREB intensity within the nuclear ROI was measured. Arc TXN intensity was calculated by dividing the intensity of the Arc transcription site by the average intensity of individual Arc mRNAs in the nucleus.

## Acknowledgments

This work was supported by the National Research Foundation (NRF) of Korea Grant No. 2020R1A2C2007285 and the startup fund from the University of Minnesota.

## Multimedia

**Movie 1. Time-lapse movie of Arc transcription induced by TBS.** The circle is the nucleus of a neuron and the two bright spots are Arc transcription sites. The timestamp indicates the elapsed time from the onset of the stimulation. This movie corresponds to Figure 1F.

